# Principal component analysis reveals multiple consistent responses to naturalistic stimuli in children and adults

**DOI:** 10.1101/2020.05.01.073163

**Authors:** Xin Di, Bharat B. Biswal

## Abstract

Functional MRI (fMRI) study of naturalistic conditions, e.g. movie watching, usually focuses on shared responses across subjects. However, individual differences have been attracting increasing attention in search of group differences or associations with behavioral outcomes. Individual differences are typically studied by directly modeling the pair-wise intersubject correlation matrix or projecting the relations onto a single dimension. We contend that it is critical to examine whether there are one or more consistent responses underlying the whole sample, because multiple components, if exist, may undermine the intersubject relations using the previous methods. We propose to use principal component analysis (PCA) to examine the heterogeneity of brain responses across subjects and project the individual variability into higher dimensions. By analyzing an fMRI dataset of children and adults watching a cartoon movie, we showed evidence of two consistent responses in the supramarginal gyrus and other regions. While the first components in many regions represented a response pattern mostly in older children and adults, the second components mainly represented the younger children. The second components in the supramarginal network resembled a delayed version of the first PCs for 4 seconds (2 TR), indicating slower responses in the younger children than the older children and adults. The analyses highlight the importance of identifying multiple consistent responses in responses to naturalistic stimuli. This PCA-based approach could be complementary to the commonly used intersubject correlation to analyze movie watching data.

## 1. Introduction

Neuroimaging study of brain functions has observed a paradigm shift from using well-controlled experimental tasks or completely unconstrained resting-state to using more naturalistic and complex stimuli such as movies and stories (Hasson et al., 2004; Nastase et al., 2019; Sonkusare et al., 2019). Compared with the resting-state, the naturalistic condition is more confined to the inputs, which could ensure that different subjects follow similar brain states. On the other hand, the stimuli are more naturalistic than arbitrarily defined trials and tasks, and maybe more efficient to elicit higher-order brain functions. When watching or listening to the same naturalistic stimuli, different subjects tend to have similar brain responses in certain brain regions (Hasson et al., 2004), which can be examined by using intersubject correlations (Chen et al., 2016; Nastase et al., 2019).

In addition to the study of shared responses, a growing research interest has begun focusing on the individual differences of responses during naturalistic conditions (Chen et al., 2017; Finn et al., 2020). Differences in shared responses have been shown between children and adults (Cantlon and Li, 2013; Moraczewski et al., 2018; Petroni et al., 2018), during aging (Campbell et al., 2015), as well as in mental disorders, such as autism spectrum disorder (Byrge et al., 2015; Hasson et al., 2009; Salmi et al., 2013) and schizophrenia (Yang et al., 2019). Within a healthy subject group, the intersubject correlations of brain responses were also correlated with the similarities of subjective ratings of the stimuli (Jääskeläinen et al., 2016; Nummenmaa et al., 2012), and subjects’ trait-like characteristics such as paranoia (Finn et al., 2018) and cognitive style (Bacha-Trams et al., 2018).

The methods for studying the individual differences in responses to naturalistic stimuli is still being developed (Chen et al., 2017; Finn et al., 2020). For a given brain region (or voxel), each subject *i* has a response time series *x*_*i*_(t), which can be partitioned into three components (Nastase et al., 2019):

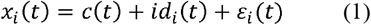

where *c*(t) represents the consistent response across subjects, *id*_*i*_(t) represents the idiosyncratic response for each subject *i*, and *ε*(t) represent noises. The idiosyncratic response *id*_*i*_(t) ideally is unique to each subject, therefore it is usually referred to as the source of individual differences. This is true by its definition. However, if everyone has different responses, it is difficult to link the responses to individual measures or group differences. In a real case scenario, it is usually assumed that there is a underlying canonical responses pattern (Finn et al., 2020), which is present across subjects but weights differently for individuals. The model can then be modified with an additional weight parameter *a*_*i*_:

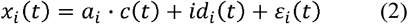

The estimates of *a*_*i*_ can then be used to correlate with group or individual differences. Here the model is different from Finn et al. (Finn et al., 2020) because they added the weight parameter to the idiosyncratic term *id(t)* rather than *c(t)*. But essentially the two models are the same because they both assume an underlying canonical response, and individual differences arise from the weightings.

A straightforward way to obtain *a*_*i*_ for is to first estimate the consistent component *c*(t), and correlate each subject’s time series *x*_*i*_(t) with *c*(t). *c*(t) is usually calculated by excluding the examined subject and averaging the remaining subjects to avoid bias, a strategy also known as leave-one-out (LOO) (Nastase et al., 2019). An alternative strategy is to calculated a pairwise intersubject correlations matrix, which projects *a*_*i*_ into two dimensions. The differences between pairs of subjects can be compared by using linear mixed-effect modeling (Chen et al., 2017). Moreover, a few models have been proposed to translate individuals’ behavioral measures into pair-wise relationships, e.g. Nearest Neighbors model and *Anna Karenina* model (Finn et al., 2020). The pair-wise relations in brain activity measures and those in behavioral measures can then be correlated to verify which model can best describe the intersubject relationships.

There are two main limitations in the current methods. First, the representational similarity approach depends on the hypothesis of the relationships. For example, *Anna Karenina* model assumes that the subjects with higher scores of a behavioral measure tend to have similar responses, but those with lower scores all respond differently. A model may not be appropriate for certain domains, and may not be able to capture complex relationships such as a non-monotonic developmental curve. Secondly, it is usually implicitly assumed that there is only one consistent component. But this may not be true in a real case scenario. For example, children may comprehend a cartoon movie differently from adults, or males and females may pay attention to different scenes and objects. Therefore, we may need to assume multiple consistent components among all the subjects. Equation 2 can then be expanded to include two consistent components *c*_*1*_(t) and *c*_*2*_(t):

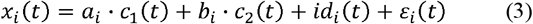

Now there are two sets of parameters to represent individual differences, *a*_*i*_ and *b*_*i*_. LOO correlation method cannot recover both sets of the parameters. And the pair-wise correlation matrix may also be difficult to capture using simple models such as *Anna Karenina* model.

We take development as an example. A certain function may start to develop after a certain age and then reach a plateau. If the function requires a certain pattern of brain responses, then the weight parameters *a*_*i*_ for that response will look like Figure 1A. A developmental curve may also like Figure 1B, where the likelihood to respond to a certain pattern first increases and then decreases as age increases. Figure 1D and 1E show the pair-wise intersubject correlations for the two developmental curves. Matrix 1D can be described by *Anna Karenina* model. But new models are needed to describe the relationships in matrix 1E. Alternatively, we can calculate LOO intersubject correlations, and the individual LOO correlations can reflect the hypothetical developmental trends (Figure 1G and 1H). A more complex scenarios is that the two consistent components may both exist and they are independent (Figure 1C).

**Figure 1.**
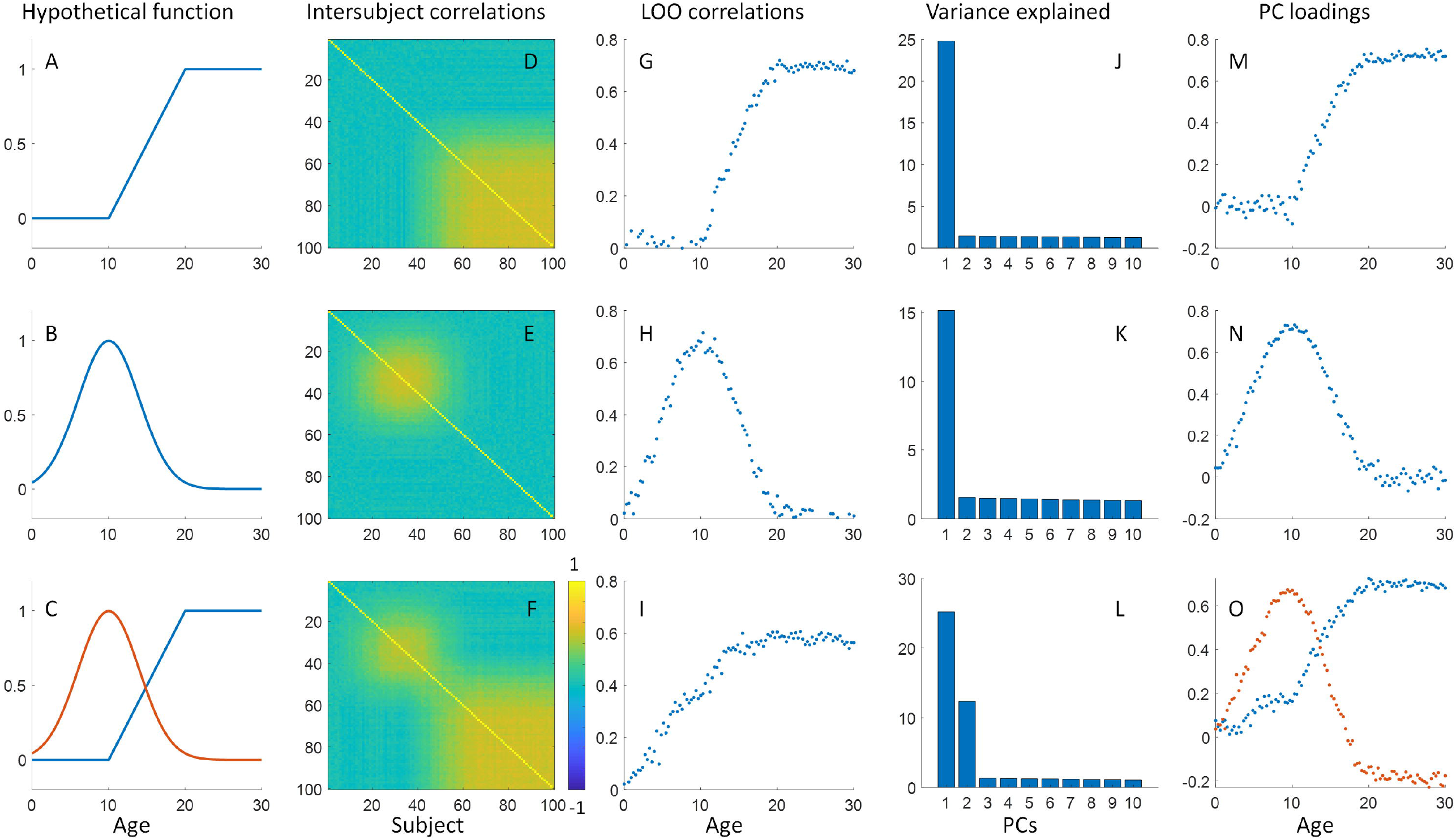
Illustrations of developmental effects of shared responses in a brain region. A and B illustrate two hypothetical developmental functions of consistent responses. The age range was set between 0 and 30 years, which overlaps with the empirical data. Note that the consistent responses in A and B may be independent. C shows the scenario where the two separate consistent components are present. D through F show the pair-wise correlation matrices across subjects. G through I show the intersubject correlations calculated using the leave-one-out (LOO) method against the subjects’ age. J through L show the percentage of variances explained by the first 10 principal components (PCs) from principal component analysis (PCA). M through O show the PC loadings of the first one or two PCs against age.

The pair-wise correlations become more complicated to be modeled (Figure 1F). And the LOO correlations can only show an averaged age effect, but cannot recover the two separate trends (Figure 1I).

To untangle the complex intersubject relationships, we proposed a principal component analysis (PCA) based analysis strategy. The time series of all the subjects form a matrix *X* (time points × subject). PCA identifies a transformation matrix *W* to transform the individual response matrix *X* into a series of principal components (PCs) *T*:

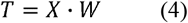

The PCs are all orthogonal. The first PC explains the largest variance of the data, and the remaining PCs similarly maximize the explained variance of the remaining variance. The variance explained by each PC is indexed by the eigenvalue of the covariance matrix of *X*, which could be an indicator of whether there are multiple consistent response patterns. In the first two hypothetical developmental functions, a single component explained a large portion of variance (Figure 1J and 1K). In contrast, in the third case with two consistent responses, the first two components both explained large portions of variance (Figure 1L).

Usually, we are only interested in the first few PCs. For the *i*^*th*^ PC, its relations to individuals’ time series are as follow:

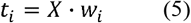

If we assume the first or second PCs as consistent responses, then the weight vector *w*_*i*_ may reflect the weight of each subject on the consistent responses. We calculated the loadings of the first PCs for the first two hypothetical developmental functions, and the loadings showed similar patterns as the developmental functions and the individual LOO correlations (Figure 1M and 1N). Moreover, the loadings of the first two PCs from the third scenario can differentiate the developmental patterns of the two consistent components (Figure 1O).

The study of group differences, e.g. a case-control study, also faces a similar problem. For example, we may expect that a group of subjects with a mental disorder have lower intersubject correlations. On the other hand, all healthy subjects might have consistent responses. But the critical question becomes whether the patient group has diminished responses at all or has a different canonical response from those in the healthy group. One can compare pair-wise intersubject correlations between groups to answer this question (Chen et al., 2017, 2016). But it could be overlooked if one only used the LOO-based method.

In summary, we have briefly reviewed the methods for studying individual differences in response to naturalistic stimuli. We argue that it is critical to examine whether there are multiple consistent components. We therefore propose a PCA-based approach to first indicate whether there are potentially multiple consistent components, and then examine individual loadings of these components. Age and biological sex are two common factors that affect the individual differences in brain structures and functions (Dosenbach et al., 2010; Lenroot et al., 2007). By using the movie-watching paradigm, two previous studies have shown higher intersubject correlations in adults compared with children (Cantlon and Li, 2013; Moraczewski et al., 2018). In the current study, by analyzing another publicly available fMRI dataset of children and adults who watched an animated movie, we ask whether there are multiple consistent response patterns in the sample. We applied our PCA-based approach and showed evidence of second consistent components in certain brain regions. We also examined the relations between the PCA-based approach and the commonly used LOO correlations in the real fMRI data.

## 2. Materials and methods

### 2.1. Data and task

The fMRI data were obtained through openneuro (https://openneuro.org/), with accession #: ds000228. There are in total of 155 subjects, with 33 adults (18 to 39 years old) and 122 children subjects (3 to 12 years old). We adopted the same criteria to remove data with poor spatial coverage and large head motion (see below) as our previous paper with only adult subjects analyzed (Di and Biswal, 2020). As a result, the adult group included 17 females and 12 males. The age range was 18 to 39 years old (*mean = 24*.*6, standard deviation = 5*.*3*). The children group included 28 females and 25 males. The age range was 3.5 to 12.3 years old (*mean = 7*.*0, standard deviation = 2*.*5*). The original study was approved by the Committee on the Use of Humans as Experimental Subjects (COUHES) at the Massachusetts Institute of Technology.

During the fMRI scan, the subjects watched a silent version of the Pixar animated movie “Partly Cloudy”, which is 5.6 minutes long (https://www.pixar.com/partly-cloudy#partly-cloudy-1). Brain MRI images were acquired on a 3-Tesla Siemens Tim Trio scanner. Younger children were scanned using one of two 32-channel custom head coils, and older children and adults were scanned using the standard Siemens 32-channel head coil. Functional images were collected with a gradient-echo EPI sequence sensitive to blood-oxygen-level dependent (BOLD) contrast in 32 interleaved near-axial slices (EPI factor: 64; TR: 2□s, TE: 30□ms, flip angle: 90°). The subjects were recruited for different studies with slightly different voxel size and slice gaps, 1) 3.13□mm isotropic with no gap; 2) 3.13□mm isotropic with 10% gap; 3) 3□mm isotropic with 20% gap; and 4) 3□mm isotropic with 10% gap). During the preprocessing, all the functional images were resampled to 3 mm isotropic voxel size. 168 functional images were acquired for each subject, with four dummy scans collected before the real scans to allow for steady-state magnetization. T1-weighted structural images were collected in 176 interleaved sagittal slices with 1□mm isotropic voxels (GRAPPA parallel imaging, acceleration factor of 3; FOV: 256□mm). For more information on the dataset please refers to (Richardson et al., 2018).

### 2.2. FMRI data processing

#### 2.2.1. Preprocessing

FMRI data processing and analyses were performed using SPM12 (SPM: RRID:SCR_007037; https://www.fil.ion.ucl.ac.uk/spm/) and MATLAB (R2017b) scripts. A subject’s T1 weighted structural image was first segmented into gray matter, white matter, cerebrospinal fluid, and other tissue types, and was normalized into standard Montreal Neurological Institute (MNI) space. The T1 images were then skull stripped based on the segmentation results. Next, all the functional images of a subject were realigned to the first image and coregistered to the skull stripped T1 image of the same subject. Framewise displacement was calculated for the translation and rotation directions for each subject (Di and Biswal, 2015). Subjects who had maximum framewise displacement greater than 1.5 mm or 1.5° were discarded from further analysis. The functional images were then normalized to MNI space using the parameters obtained from the segmentation step with a resampled voxel size of 3 × 3 × 3 mm^3^. The functional images were then spatially smoothed using a Gaussian kernel of 8 mm. Lastly, a voxel-wise general linear model (GLM) was built for each subject to model head motion effects (Friston’s 24-parameter model) (Friston et al., 1996), low-frequency drift via a discrete cosine basis set (1/128 Hz cutoff), and a constant offset. The residuals of the GLM were saved as a 4-D image series, which were used for further intersubject correlation analysis.

Regarding the potential head motion effects, we firstly calculated framewise displacement in translation and rotation separately (Di and Biswal, 2015), and removed subjects with maximum framewise displacement greater than 1.5 mm or 1.5°. As a result, 82 subjects were included in the final analysis. Secondly, we performed PCA on the framewise displacement time series in translation and rotation. The first components explained 4.66% and 4.30% of the variance in the two directions, respectively, suggesting that there were very limited intersubject correlations of head movements across subjects (also see Supplementary Figure S1). Thirdly, in preprocessing 24 head motion variables have been removed from the fMRI time series. Lastly, we calculated mean framewise displacement in translation and rotation for each subject. The children group showed significant larger mean framewise displacement in rotation compared with the adult group (*t = 6*.*04, p < 0*.*001*) (see Supplementary section S1 for details). In later analyses considering age effects or behavioral correlations, we regressed out the mean framewise displacements in translation and rotation from the PC loadings and compared the results before and after the regression.

### 2.2.2 Dimension reduction

We first focused on a small number of large-scale networks, which enabled us to perform an in-depth analysis of their time courses and individual variations. We performed spatial independent component analysis (ICA) to define large-scale networks by using Group ICA of fMRI Toolbox (GIFT: RRID:SCR_001953; http://mialab.mrn.org/software/gift) (Calhoun et al., 2001). Twenty components were extracted. The resulting IC maps were visually inspected, and fifteen maps were included in the subsequent analysis as functionally meaningful brain networks. The full maps of all the 20 ICs can be found at: https://neurovault.org/collections/INSJUAIW/. For each IC, a time series was back-reconstructed to each subject using the group ICA method, resulting in a 168 (time points) × 82 (subject) matrix. To avoid confusion with PCA in the current paper, we refer to the IC maps as networks below. Secondly, we performed PCA on a voxel basis to study the spatial distributions.

### 2.3. Principal component analysis

For each network (IC) or voxel, we performed PCA on a 168 (time points) × 82 (subject) matrix *X*. The time series of each subject (each column) was first z transformed, which is a critical step in PCA. Then, PCA was performed in MATLAB by using the singular value decomposition algorithm. The eigenvalues of the covariance matrix of *X* were obtained. The percentage variance explained by each PC was then be calculated as the corresponding eigenvalue divided by the sum of all the eigenvalues. The PC scores (time series) and the associated weighting for each subject *w*_*i*_ were also obtained. PC loadings were calculated as the PC weights multiplying the standard deviation of the eigenvalue.

To determine whether a PC explained greater variance than the random level, we performed a circular time-shift randomization to determine the null distributions (Kauppi et al., 2010). The time-shift method can preserve the autocorrelations in the BOLD time series, which is preferable to a simple permutation test. Specifically, each subject’s time series were added a delay drawn from a discrete uniform distribution of 0 to 167 with replacement, then the PCA was performed, and the variances explained by the first PC was obtained. The process was repeated 10,000 times to form a null distribution. The variances explained by the first two PCs from the real fMRI data were compared with the null distributions to obtain the p values. It is noteworthy that the null distributions were calculated based on the first PC, which is a conservative choice for the statistics of the second PCs.

For the ICA-based analysis, we performed the circular time-shift randomizations for every network (ICs). We adopted a threshold of p < 0.001 to account for the multiple comparisons. An alternative approach is to use false discovery rate (FDR) correction. However, FDR depends on the overall distributions of all the regions. It may make the thresholding different among different spatial scales. We therefore adopt the same threshold of p < 0.001 for the ICA-based and voxel-wise analyses, which was more stringent than FDR corrected p < 0.05 in the current case. The randomization was quite computationally expensive for the voxel-wise analysis. Therefore, we performed PCA on 1,000 regions (Schaefer et al., 2018) and calculated the local null distributions. The voxel-wise PCA results were compared with the null distribution in a local region to compute the p values.

Because the later PCs may represent only a small number of subjects, we evaluated whether the PCs could be reliability identified with sample variations. We performed bootstrapping along the subject dimension for 1,000 times. PCA was performed on the bootstrapping samples, and the correlations of the PCs were calculated among the samples. The goal is to verify whether the identified PCs were consistent across the bootstrapping samples. The 95% confidence interval of the variance explained by the second PCs were also obtained. This analysis was only performed for the ICA-based analysis.

### 2.4. Cross-correlation and delay estimates

Because we found that the second PC score in some networks seemed to be delayed to the first PC, we calculated cross-correlations between the two PC scores to confirm this. The autocorrelation in BOLD signals could produce spurious cross-correlations (Dean and Dunsmuir, 2016), therefore, we performed simulations with components of convolution with hemodynamic response function (HRF). Specifically, we generated two Gaussian time series with 168 time points and convolved them with the canonical HRF in SPM. Cross-correlations were then calculated, and the maximum absolute value of the cross-correlations was obtained. The procedure was repeated 100,000 times to form a null distribution of the maximum value. The 95 percentile of the distribution was used as the critical value for the cross-correlation analysis for the real fMRI data.

We also calculated the time lags between the time series of every subject with reference to the first PC score by obtaining the time point of maximum absolute cross-correlation. Because single subject time series were noisy, we set a maximum lag of ±5 time points in search of lags.

### 2.5. Behavioral correlates

We next asked whether the first two PCs of different networks can provide complementary information in explaining the variability of a behavioral measure. Test scores of theory of mind performance are available for the children subjects (n = 53). The theory of mind battery includes custom-made stories and questions that require an understanding of the characters’ mental states. The theory of mind task performance was summarized as the proportion of correct questions out of the 24 items. More information about the task and scores can be found in Richardson et al. (2018). The analysis was only performed for the ICA-based analysis, where the second PC explained significant variance (i.e. the supramarginal network). We first examined simple correlations between the first or second PC loadings and the theory of mind performance. Next, we put the two PC loadings together in a linear regression model to explain the variance of the theory of mind performance. The t statistics corresponding to the two PC loading regressors were obtained.

### 2.6. LOO correlation and the relations to PCA-based method

Although the main focus of this study is to apply PCA to identify potentially multiple consistent responses, PCA can also provide measures on intersubject correlations. Specifically, we asked whether the variance explained by the first PC is related to the averaged intersubject correlations and whether the loadings of PC1 are related to the individual LOO correlations. For a specific region, we calculated LOO intersubject correlations on the 168 (time points) × 82 (subject) matrix *X*. Specifically, a subject’s time series was held out and the consistent component was calculated by averaging the remaining 81 subjects. Then the correlation between the subject’s time series and the averaged time series was calculated. Each subject then had a LOO correlation value. The LOO correlations were Fisher’s z transformed, averaged, and then transformed back to r values to form an averaged intersubject correlation in a region.

We first examine the relationships on all the 20 networks (ICs). The averaged intersubject correlations were squared to match with the variance quantity. We then calculated the correlations between the variance explained by the first PC and the squared mean correlations across the 20 networks. Next, for each network, we calculated the correlations between the first PC loadings and individual LOO correlations. The same analysis was performed on the 1,000 ROIs.

The rationale for including the noise ICs in the analysis is to reveal more general relations between PCA and LOO correlations. Imagine if all the time series are noise, given the high dimensionality (number of subjects), then the first PC may not be identified as the averaged time series. But if there are underlying consistent signals, then the first PC may turn out to be very similar to the averaged time series, i.e. the consistent response. This will in turn give rise to high correlations between PC1 loadings and individual LOO correlations. We performed a simple simulation to reveal such a relationship. We generated a 168 × 82 matrix with a 168 Gaussian vector representing the consistent response and a 168 × 82 Gaussian random matrix representing the noises. The consistent component had different weights for subjects drawn from a uniform distribution between 0 and 1. And finally, the subjects’ weights were multiplied by an overall weight value from a uniform random distribution (from 0 to 1) to vary the overall levels of intersubject correlations. The procedure was repeated 1,000 times. We then calculated PCA and LOO correlations and examined their relations.

## 3. Results

### 3.1. ICA-based analysis

We first performed PCA on the 15 large-scale networks and obtained the percentage of variances explained by the PCs (Figure 2). The first PCs of all the 15 networks explained more than chance-level variance at p < 0.001. Among the 6 networks that explained the highest variance (more than 15%), five were visual related networks and the remaining IC 17 was located in the supramarginal gyrus. These are consistent with our previous voxel-wise analysis in only adult subjects (Di and Biswal, 2020). For the second PCs, only the supramarginal network (IC17) explained more than chance-level variance (*6*.*01%, p < 0*.*001*). Supplementary Figure S4 shows the variance explained by all the PCs in this network. It is noteworthy that the variance explained by the second PC would be much less than those explained by the first one. But it may be still meaningful, because the differences may reflect the number of subjects represented in different PCs. To evaluate the stability of the PC2, we performed a bootstrapping along the subject dimension. Supplementary Figure S5A and S5B show that PC2 could be reliability identified among the bootstrapping samples. And the 95% confidence interval of the explained variance by PC2 was between *5*.*65%* and *7*.*50%*.

**Figure 2.**
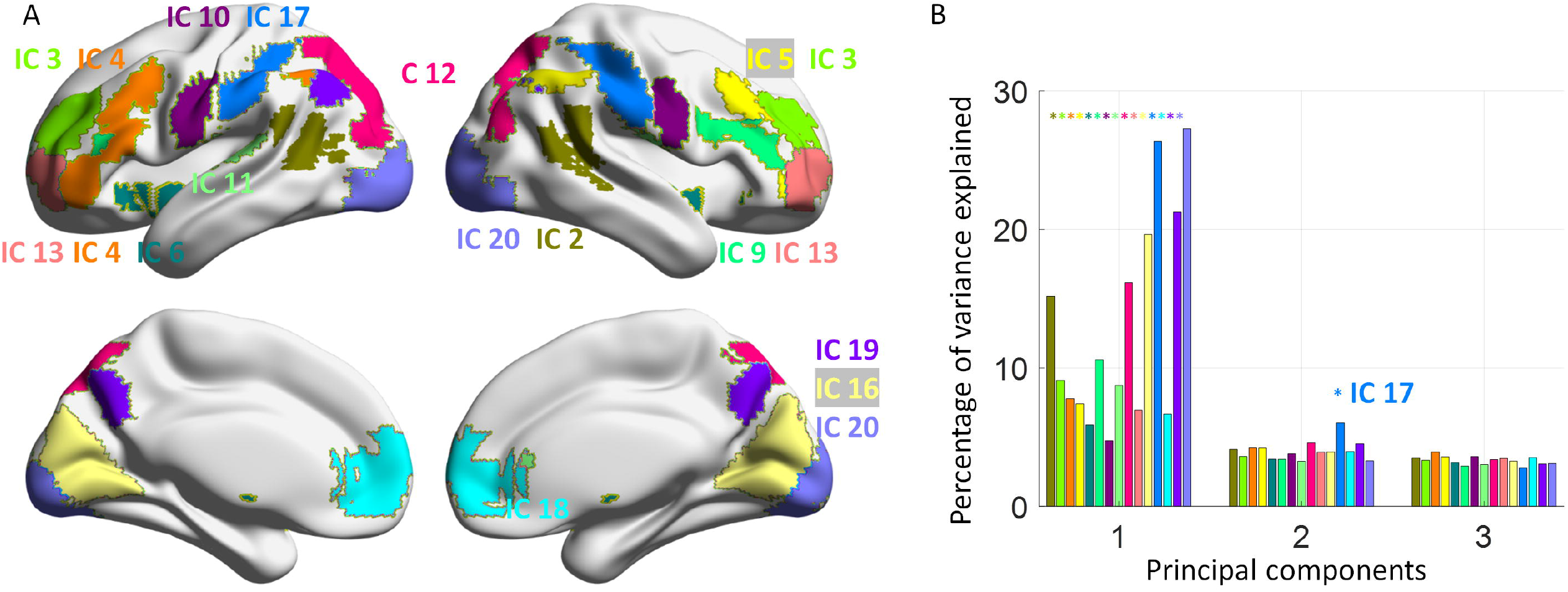
A. Maps of 15 independent components (ICs) that are included in the current analysis. The group averaged maps were thresholded at z > 2.3. B. Percentage of variance explained by the first three principal components for the 15 networks (ICs). The bar colors correspond to the network colors in panel A. * represents p < 0.001 by using a circular time-shift randomization method. The brain networks were visualized with BrainNet Viewer (RRID: SCR_009446) (Xia et al., 2013).

The pair-wise correlation matrix in the supramarginal network clearly showed a trend of greater intersubject correlations in older subjects (Figure 3A). The PC1 loadings were mostly positive and were greater as age increased and reached a plateau during the adult age range. In contrast, the PC2 loadings were positive for the younger children but mostly negative for the adults. No clear sex differences can be found in both PCs. We also calculated the LOO intersubject correlations (Figure 3D), which turned out to be very similar to the PC1 loadings (*r* ≈ *1, p < 0*.*001*, see also the scatter plot in supplementary Figure S6). The mean framewise displacement in rotation showed small but statistically significant correlations with both PC1 loading (*r = -0*.*30, p = 0*.*007*) and PC2 loading (*r = 0*.*29, p = 0*.*009*). We therefore regressed out the mean framewise displacements from the two PC loadings. The age effects on the adjusted PC loadings remain very similar to what on the original PC loadings (Figure S2).

**Figure 3.**
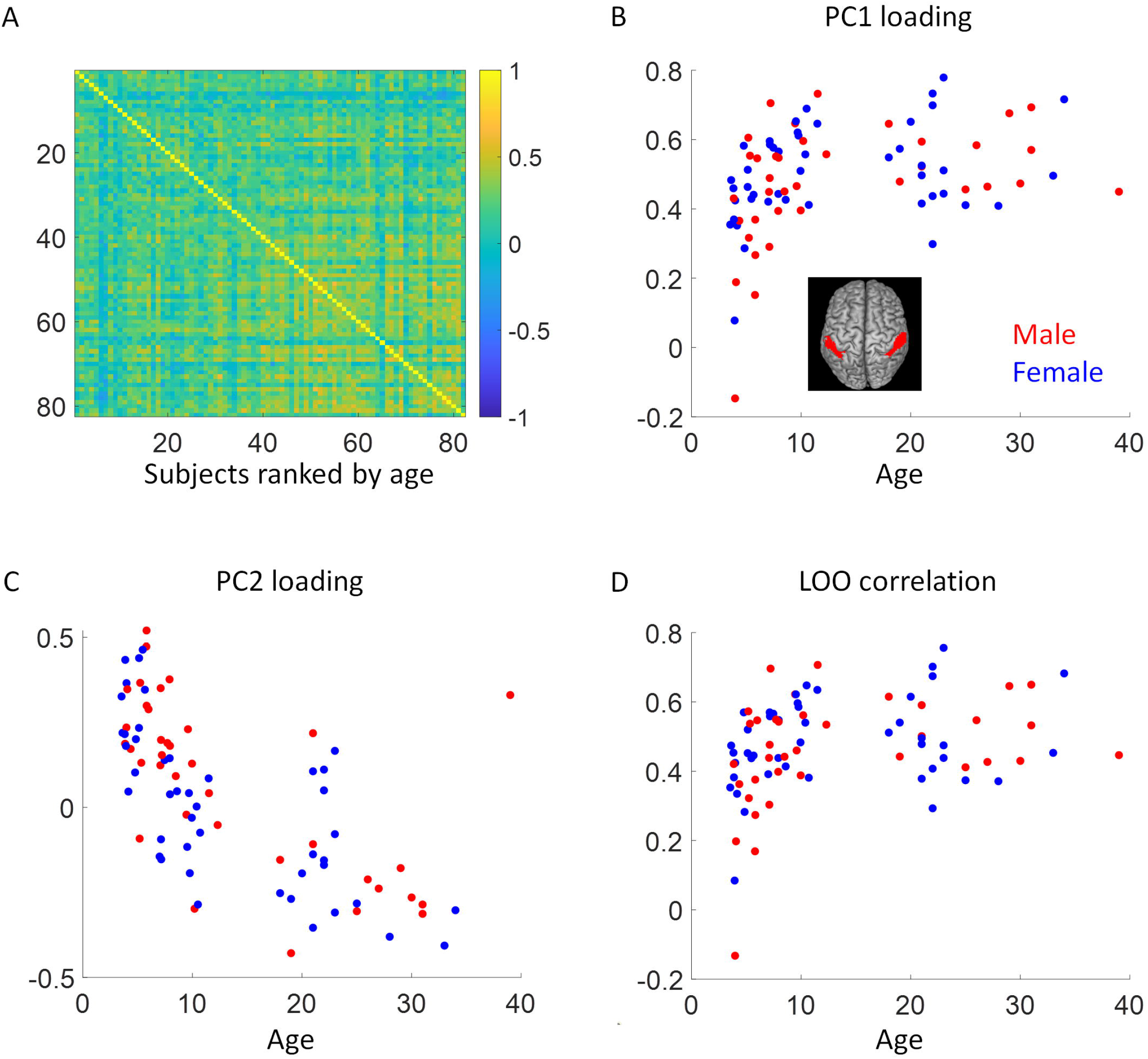
A, Correlation matrix of the supramarginal gyrus network (independent component 17) across the 82 subjects. The subject were sorted by age in an ascending order. B and C, Principal component (PC) loadings for the first and second PCs as functions of age. D, Leave-one-out (LOO) intersubject correlations as a function of age. The brain slice illustrates the location of the network.

Figure 4A shows the time series of the first two PCs (PC scores) in the supramarginal network. Interestingly, PC2 looked similar to PC1 but seemed delayed compared with the PC1. Cross-correlation analysis confirmed a 2-TR (4 s) delay between them (Figure 4B). Further, we examined whether the loadings of the PC2 reflect the lags of an individual’s time series. We calculated the time shifts between each individual’s time series with reference to the PC1 time series. 80 out of the 82 subjects had a -1 to 1 time points shifts. The individual’s time shifts related to PC1 were highly correlated with the PC2 loadings (Figure 4C). In Supplementary Figure S8, we show individual time series with subjects ordered according to age (top row) and the PC2 loadings (bottom row). It shows clearly that for older subjects the time series appeared to be faster compared with the younger subjects. The time lags became clearer when the subjects were sorted by the PC2 loadings.

**Figure 4.**
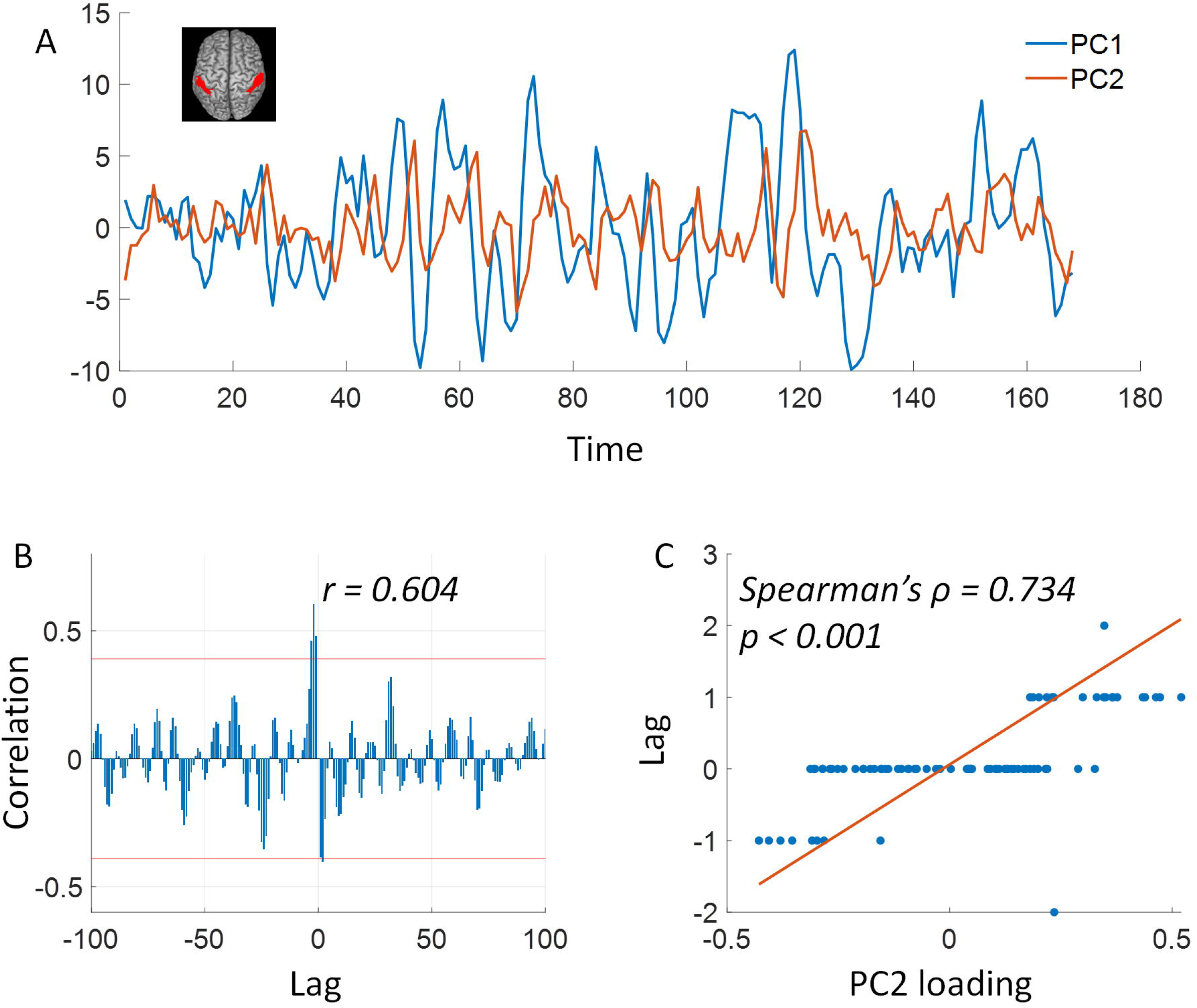
A, Principal component (PC) scores of the first two PCs in the supramarginal network (independent component 17). The brain slice illustrates the location of the network. B, Cross-correlations between the first two PCs. The red lines indicate p < 0.05 of absolute peak cross-correlations. C, Time shifts of individual’s time series with reference to the first PC score as a function of the PC2 loading.

We next asked whether the two PCs of the supramarginal network (IC17) can provide complementary information in explaining the variations of the theory of mind task performance (proportion correct). The first PC loadings were positively correlated with the theory of mind performance (*r = 0*.*468, p < 0*.*001*, Figure 5A), and the second PC loading was negatively correlated with the theory of mind performance (*r = -0*.*398, p = 0*.*003*, Figure 5B). However, when including both the first two PC loadings in a linear regression model to predict the theory of mind performance, only the first PC loadings had a statistical significant effect (*t*_*PC1*_ *= 2*.*47, p*_*PC1*_ *= 0*.*017*; *t*_*PC2*_ *= -1*.*44, p*_*PC2*_ *= 0*.*157*). This is probably due to the fact that the two PC loadings were correlated (*r = -0*.*532, p < 0*.*001*). We also regressed out the mean framewise displacement in both translation and rotation from the PC loadings. The correlations between the adjusted PC loadings and the theory of mind performances remained significant (PC1: *r = 0*.*428, p = 0*.*001*; PC2: *r = -0*.*278, p = 0*.*044*). Lastly, the LOO intersubject correlations were also correlated with theory of mind performance (*r = 0*.*461, p < 0*.*001*).

**Figure 5.**
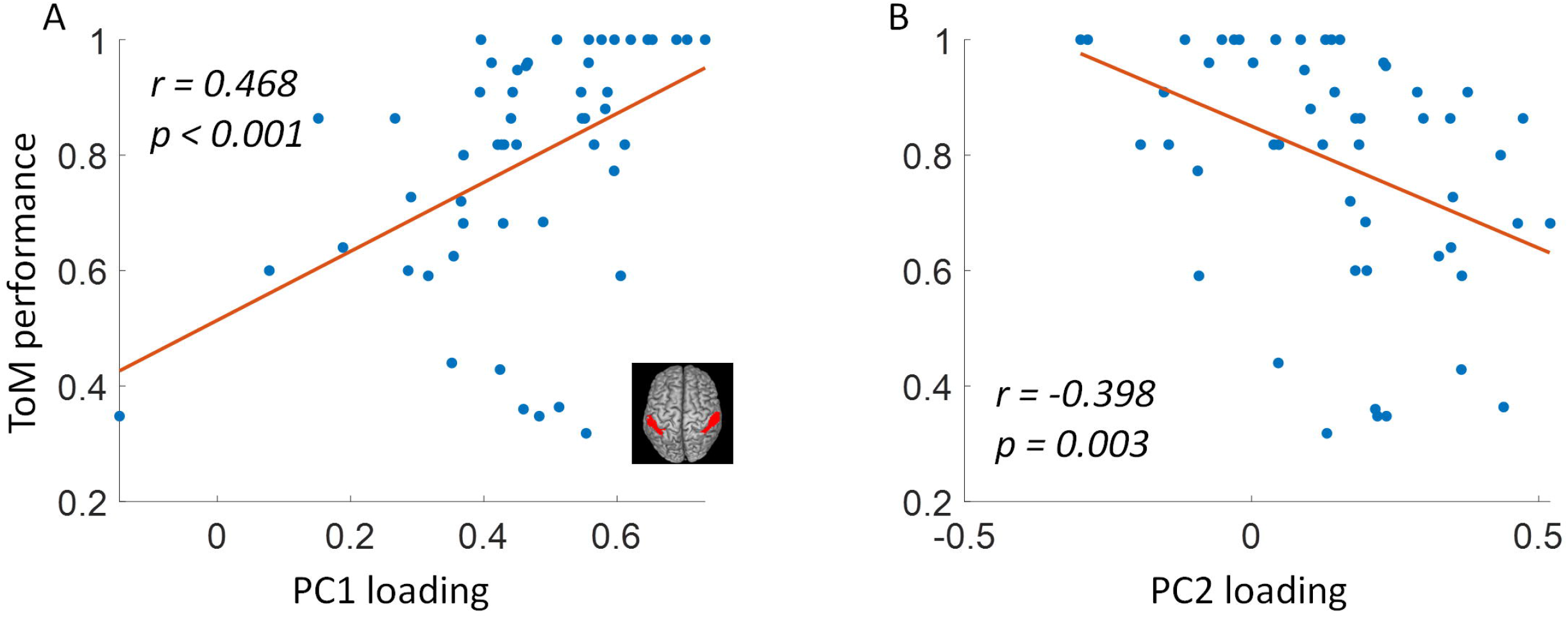
Correlations between theory of mind (ToM) performance (proportion of correct) and principal component (PC) loadings for the first two PCs of the supramarginal gyrus network (independent component 17). The brain slice illustrates the location of the network.

### 3.2. Spatial distributions of variances explained by the second PCs

The left panel of Figure 6 shows the spatial distributions of significant second PCs at p < 0.001. For reference, the PC1 variance map is shown in the right panel. One major cluster of the PC2 map covered the supramarginal gyrus and extended to the posterior parietal lobe and posterior visual regions. We further increased the threshold to 5% to break it into three small clusters, including two clusters covering the left and right supramarginal gyrus and one cluster in posterior visual areas. We extracted the averaged time series in these clusters and performed PCA. The first two PC loadings in the three clusters were very similar to those of the supramarginal network (IC 17), i.e. the first PC loadings reflected a maturation age effect and the second PC loadings reflected higher weights in younger children.

**Figure 6.**
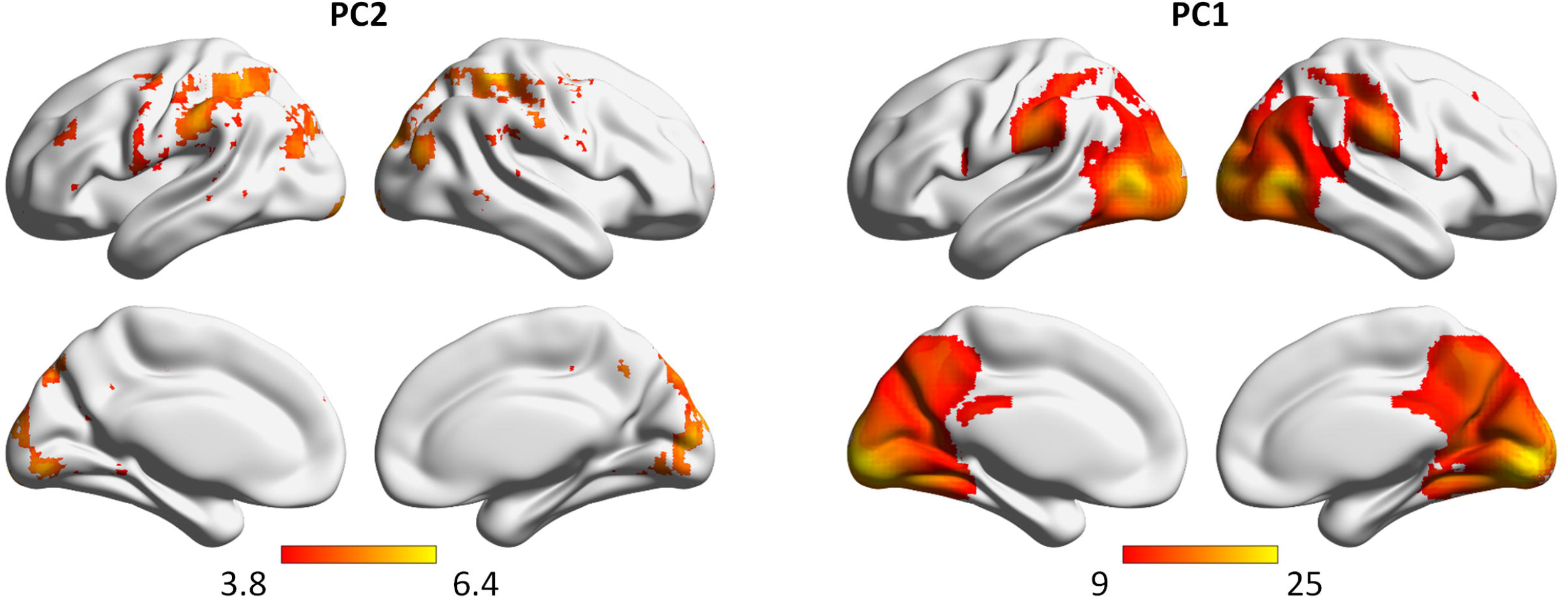
Percentage of variance explained by the second (A) and first (B) principal components (PCs) from the voxel-wise analysis (C). The voxels in A were thresholded at p < 0.001. The voxels in B were thresholded at 9%. The brain networks were visualized with BrainNet Viewer (RRID: SCR_009446) (Xia et al., 2013).

Outside the major cluster, there were three clusters larger than 40 voxels at p < 0.001, including the precuneus and left and right sensorimotor regions. The preceneus, which is part of the default mode network, is particular interesting given its role in theory of mind processing (Richardson et al., 2018).

The first two PCs and their loadings of the precuneus are shown in Figure 7. Similar to the supramarginal network (IC 17), the first PC loadings showed a maturation age effect and the second PC loadings had higher weights in younger children. The PC2 time series also seemed to be a delayed version of PC1 but with a 2-TR (4 s) lag (Figure 7A and 7D).

**Figure 7.**
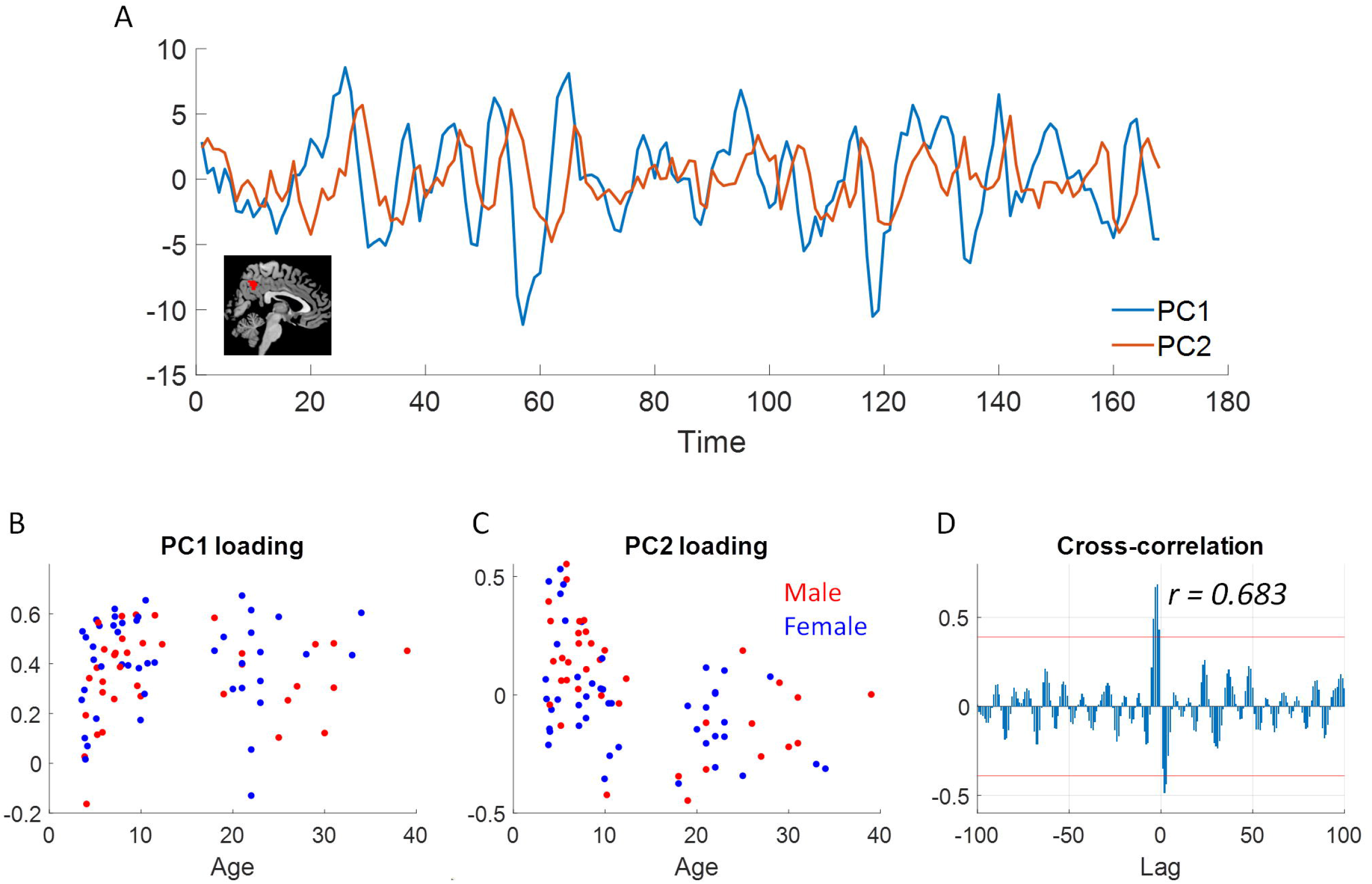
A, the time series of the first two principal components (PC scores) for the precuneus region (depicted in the brain slice). B and C, the first and second principal component (PC) loadings as functions of age. D, the cross-correlations between the two PCs. The red lines indicate p < 0.05 of absolute peak cross-correlations.

The left and right sensorimotor regions had very similar time courses and age effects. Figure 8 shows the left sensorimotor region as an example. In contrast to the previous networks and regions, the PC1 loadings of the left sensorimotor region first increased with age in the children group, but decreased with age in the adult group. Conversely, the PC2 loadings had higher weights in the adult group. The cross-correlation between PC1 and PC2 also had maximum correlation at 2-TR (4 s) lag, but PC2 was 2-TR in advance. Because PC1 had large weights in younger subjects and PC2 had larger weights in older subjects and PC, the cross-correlation indicated that the older group had a faster brain activity compared with the younger group, which is consistent with the previous networks and regions. Lastly, the age effects of the PC loadings were not confounded by the head motions. When regressing out framewise displacement from the PC loadings, the age effects remained very similar (Supplementary Figure S3).

**Figure 8.**
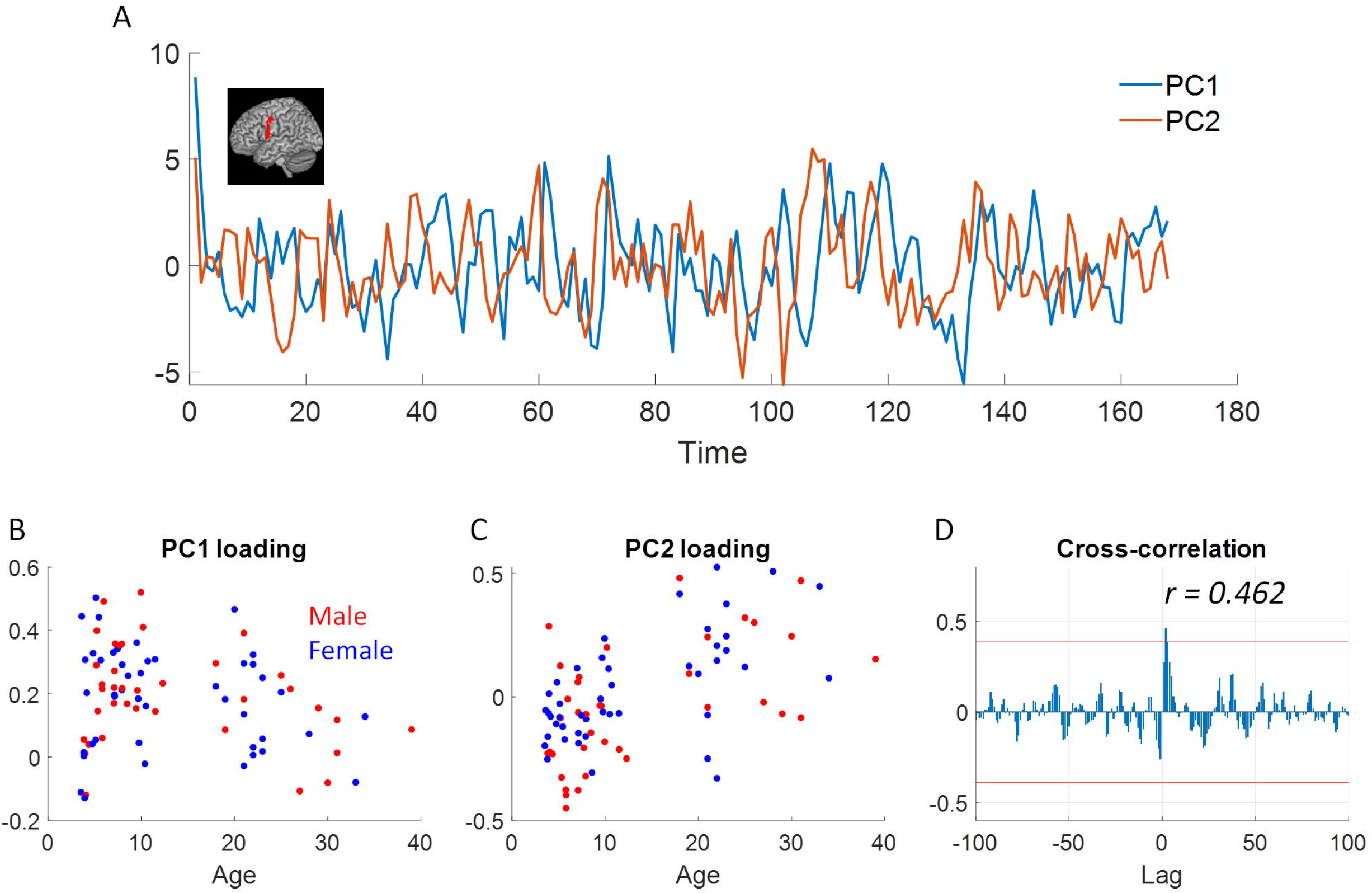
A, the time series of the first two principal components (PC scores) for the left sensorimotor region (depicted in the brain slice). B and C, the first and second principal component (PC) loadings as functions of age. D, the cross-correlations between the two PCs. The red lines indicate p < 0.05 of absolute peak cross-correlations.

### 3.3. Relations to LOO correlations

Lastly, we asked whether the PCA-based measures are related to the commonly used intersubject correlation measures. Among the 20 networks (ICs) and the 1,000 ROIs, we found almost perfect linear relations between the squared mean intersubject correlations and the percentage of variance explained by the first PCs (Figure 9A and 9B). However, their relations were off the *y = x%* line, suggesting that their quantities were not directly comparable. A similar relation can be found in the simulations (Figure 9C).

**Figure 9.**
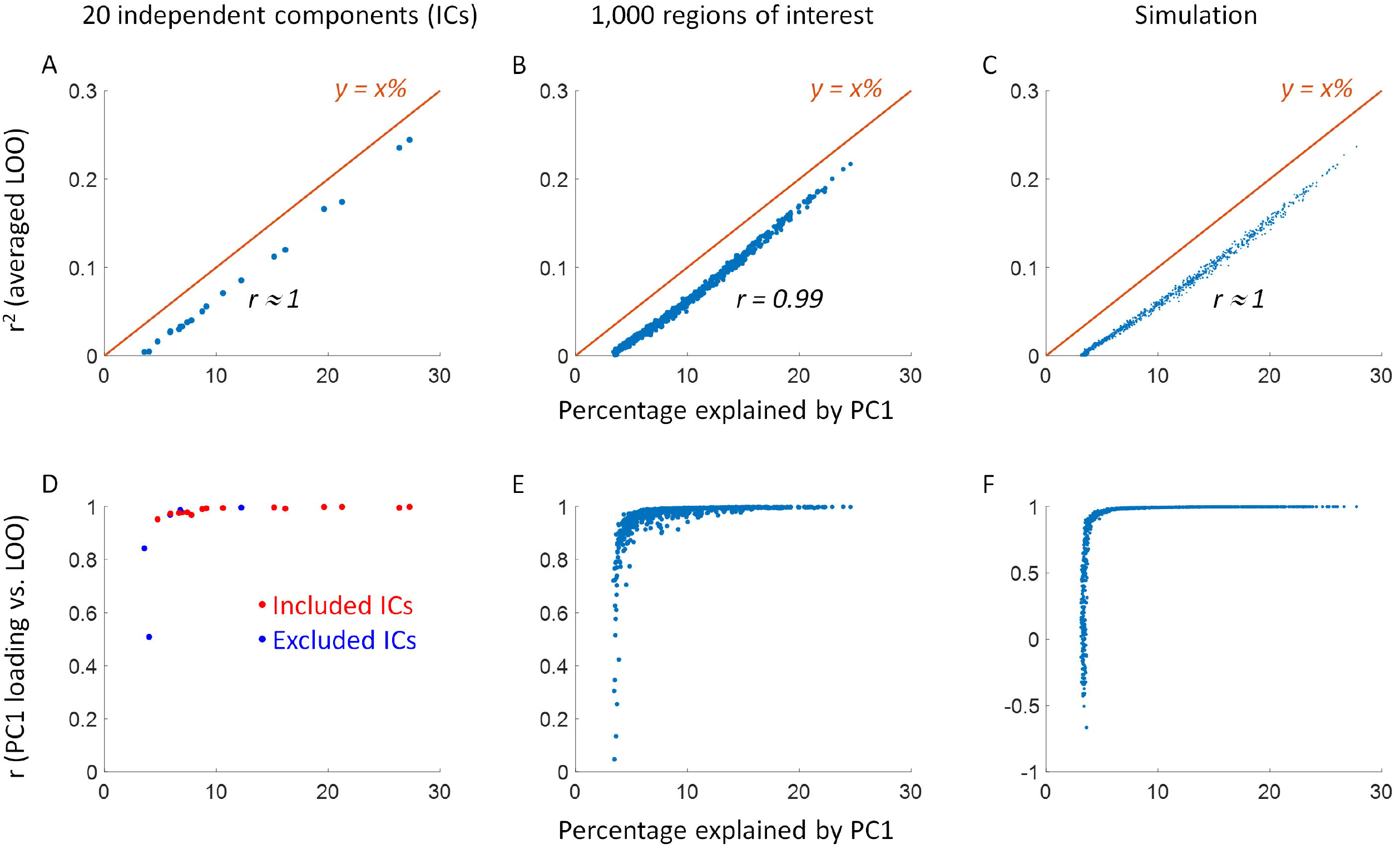
Upper row, correlations between the squared mean intersubject correlations using leave-one-out (LOO) method and the variance explained by the first principal component (PC). Lower row, the correlations between the first PC loadings and individual LOO correlations as functions of the variance explained by the first PC.

As expected, we found that the correlations between PC1 loadings and individual LOO correlations are related to the noise level of a region. We therefore plotted the correlations against the variance explained by the first PC in a network (Figure 9D) and ROI (Figure 9E). We found that if the variance explained by the first PC were higher than 5%, i.e. there are likely underlying consistent responses, then the correlations between the PC1 loadings and individual LOO correlations were higher than 0.95. But if there were very low variance explained by the first PC, then the correlation could drop to 0.5. The two networks with low correlations in Figure 9D were both considered noise components, which were excluded in the current analysis. Such a relationship was confirmed by the simulation data (Figure 9F). The simulation results further showed that in noisy conditions the correlations between PC1 loadings and individual LOO correlations varied in a wide range. But if there were underlying consistent signals, then their correlations could be close to 1. In supplementary Figure S9, we further show that the correlations between PC1 loadings and LOO correlations were related to whether PC1 could capture the averaged signals. In other words, the higher the correlations between PC1 and averaged signals, the higher the correlations between PC1 loadings and LOO correlations.

## 4. Discussion

In the current paper, we proposed a PCA-based framework to study the individual differences in response to naturalistic stimuli in fMRI data. On a movie watching dataset of children and adults, we showed evidence of second PCs in distributed brain regions, which may represent a second consistent response to the movie in the tested sample. The two PCs showed different distributions in age but not in biological sex, suggesting that the two consistent responses represent different age groups. The regions that showed the second consistent responses were in the supramarginal gyrus, posterior parietal lobe, visual areas, the precuneus, and sensorimotor regions. Interestingly, in the supramarginal gyrus, the second PC represented delayed responses than the first PC for 4 seconds (2 TR), suggesting the children around 5 years old may have delayed response compared with the adults. The results indicate the importance of studying potentially multiple consistent responses in large samples.

By calculating the eigenvalues of the covariance matrix, we provided evidence of potentially multiple consistent responses in the sample, which cannot be identified by using intersubject correlations. It is noteworthy that the variance explained by the second PCs in the current study were around 5% to 6%, which were much smaller than those by the first PCs. It may reflect the fact that the second PCs only represented a small number of subjects, but not that the correlations among them were lower. By using bootstrapping on the subject dimension, we showed that the second PC in the supramarginal gyrus were reliable against subject sampling, which support the former interpretation. As the sample sizes in neuroimaging studies become larger and larger, it becomes more important to identify sub-groups of subjects with distinct but consistent responses from other subjects. PCA provides an unsupervised tool to visualize and identify the potential sub-groups.

The regions that showed evidence of a second consistent components included the supramarginal gyrus, the posterior parietal lobe, higher visual areas, the precuneus, and sensorimotor regions. Except for the sensorimotor regions, the other regions seemed to follow similar subject weightings. That is, the first PC represented an increasingly similar response as age increased, and the second PC represented a higher similar response in children around age 5. These all suggest that the children around 5 years old showed a unique pattern of brain responses compared with both the adults and the other children groups.

The supramarginal network is a critical region involving in the theory of mind process (Silani et al., 2013) and the understandings of others’ pain (Bruneau et al., 2015). In the current results, the loadings of both PC1 and PC2 were correlated with the theory of mind performance, further confirmed its role in the understanding of the movie. Interestingly, we found that the PC2 seemed to be a delayed version with reference to the PC1, and the PC2 loadings could reflect the delays of an individual’s time series. This suggests a multivariate nature of functional developments in this region. For the youngest children in this sample, theory of mind has not been fully developed. The functional responses in the supramarginal gyrus did not show similarity among each other, nor with the older children or adults. For the children around 5 years old, the theory of mind ability has started developing, but the brain responses may be less reliable and slower compared with adults. As growing older, the responses becomes more reliable and similar to the adults. A study using the same movie stimuli has shown that when the movie was shown the second time to children of 6-7 years old, the responses shifted earlier than those from the first presentation (Richardson and Saxe, 2020), further support that the brain response time may reflect the ability of the understanding the movie.

One consideration related to the delays in BOLD signals is the inherent autocorrelation (Friston et al., 2000, 1994). Usually, the BOLD signals have a high autocorrelation at 1-TR (2 s) lag, and remain a small autocorrelation at 2-TR (4 s) lag. This means that if the delays are with 2 seconds, then PCA probably will not able to identify two distinguished components. Moreover, PCA forces the latter PCs to be orthogonal to the former PCs, meaning the signals related to PC1 have been removed from PC2. This may make PC2 look spikier than PC1. In other words, PC2 doesn’t simply represent one particular group of subjects, but those after considering the PC1 effects. One can think of PC2 as a higher-order deviation to PC1 that captures certain individual variations in the sample.

In the voxel-wise analysis, we observed distributed regions in the posterior parietal lobe, higher visual areas, the precuneus, and sensorimotor regions who showed evidence of second consistent components. The precuneus is particularly interesting given its role in theory of mind processing (Richardson et al., 2018). The other regions may be related to attention and sensorimotor processing. Previous studies have suggested that children and adults activate different brain regions when watching real versus cartoon movies (Han et al., 2007, 2005). They found that the medial prefrontal cortex was activated in children but not adults when watching cartoon movies. Although the regions identified are different, all the studies have suggest different brain response patterns between children and adults.

PCA is an unsupervised approach that relaxes the assumptions on the interindividual relationships. It is particularly useful for continuous variables such as age, where the exact timing of developments may be unknown or the developmental effects may not be monotonic. Two previous studies have compared the pair-wise intersubject correlations between children and adults and found reduced intersubject correlations in children (Cantlon and Li, 2013; Moraczewski et al., 2018). These were done by defining specific age groups. When using the PCA-based approach or LOO-based approach (Campbell et al., 2015), age can be treated as a continuous variable, so that the age effects can be modeled as developmental trajectories. On the other hand, when using the intersubject representational similarity analysis approach (Finn et al., 2020), the age effect may be captured by *Anna Karenina* model, where only older subjects respond more similarly to each other. But this model cannot capture a non-monotonic age effect, nor different consistent responses. One may need to develop new models to capture complex age effects when using the representational similarity approach.

In addition to the information about multiple consistent components, PCA can also provide similar information as LOO intersubject correlations. The variance explained by the first PC is a similar measure as averaged intersubject correlations. The current results showed that across brain regions the variance explained by the first PC was almost perfectly correlated with the averaged intersubject correlations. Moreover, the loadings of the first PC provide a simple way to project the consistent response to an individual’s dimension, which is easier than correlating each subject’s time series with the LOO averaged time series (Nastase et al., 2019). The current results showed very high correlations between PC1 loadings and LOO intersubject correlations in real fMRI data. A simple simulation also suggested that when there were underlying consistent components, PC1 loadings and LOO correlations were highly correlated. Therefore, the PCA-based method can provide similar information as LOO intersubject correlations.

There are also limitations regarding the PCA-based method. First, the baseline of the variance explained by the first PC is not zero (the x-intercept in Figure 9A through 9C), and is related to the number of subjects. When there are *n* subjects, imagine if the first PC is randomly assigned as one subject’s time series, it will explain 1/n variance. Therefore, the variance explained by the first PC cannot be compared between different sample sizes. Second, statistical testing for the PCA related parameters are not straightforward. In the current study, we adopted randomization-based nonparametric methods, which are time-consuming.

A more general challenge in studying naturalistic stimuli is the interpretations of the observed consistent responses. It becomes more difficult when multiple consistent responses are identified. In the current data, we found that delays of the signals may explain the differences between the two PCs. There may be other factors that could contribute to the differences. Future study may need to formulate testable hypotheses regarding the brain responses in different age groups to examine the causes of the differences further. More generally, when there are multiple consistent responses, reverse correlation technique (Hasson et al., 2004; Richardson et al., 2018) could be used to identify the events represented in different consistent responses. Advanced encoding models may also be helpful to explain the underlying coding of different consistent responses (Bartels et al., 2008; Nishimoto et al., 2011). But it could be difficult when the effects of interest are higher-order social processes such as theory of mind. Secondly, studies have shown that the shared responses are dynamic (Di and Biswal, 2020; Simony et al., 2016). The presence of multiple response components and their relations may also be sensitive to the movie context, thus showing fluctuations. For example, delays in responses may only occur to certain events, but not throughout the whole time series. Further studies may take dynamics into account to fully characterize the individual differences in responses. Lastly, the current study is limited by the sample size and scan time of each subject. Further study is needed with larger sample size and longer scan time to evaluate the generalizability and reliability of the current findings.

## 5. Conclusion

When watching movies, the brain may respond similarly or idiosyncratically across individuals. It is also possible that multiple consistent responses exist in different subgroups, which is overlooked by the currently available methods. We proposed a PCA-based approach to analyze the individual differences in response to naturalistic stimuli, which can detect the potential multiple consistent responses. With an example movie watching data of children and young adults, we showed evidence of two consistent responses in many brain regions, one more weighted in the adults and the other more weighted in younger children. The results highlight the importance of identifying multiple consistent components when studying shared responses to naturalistic stimuli. And PCA could be a complementary approach to analyze naturalistic stimuli data.

## Acknowledgment

This study was supported by grants from the National Institute of Health, United States (R01 AT009829; R01 DA038895).

