## Supplementary materials for "Principal component analysis reveals multiple consistent responses to naturalistic stimuli in children and adults"

**Table of contents**

S1. Head motion effects

S2. Miscellaneous Analysis on the second PC of the supramarginal network

S3. Correlations between PC1 loadings and LOO correlations in the supramarginal network

S4. Miscellaneous Analysis on the delay effects in the supramarginal network

S5. Relations between principal component analysis and LOO correlations

**S1. Effects of head motions**

Firstly, we show that head motions were not synchronized across subjects. We calculated framewise displacements (FD) for translation and rotation separately for each subjects. Figure S1A and 1D show the intersubject correlation matrices of the FD time series in the two directions. The first components explained 4.66% and 4.30% of the variance in the two directions, respectively, suggesting that there were very limited intersubject correlations of head movements across subjects. Figure S1B and S1E show the first principal component (PC) loadings as functions of age in the two directions, respectively. The PC1 loadings did not show clear age effects.

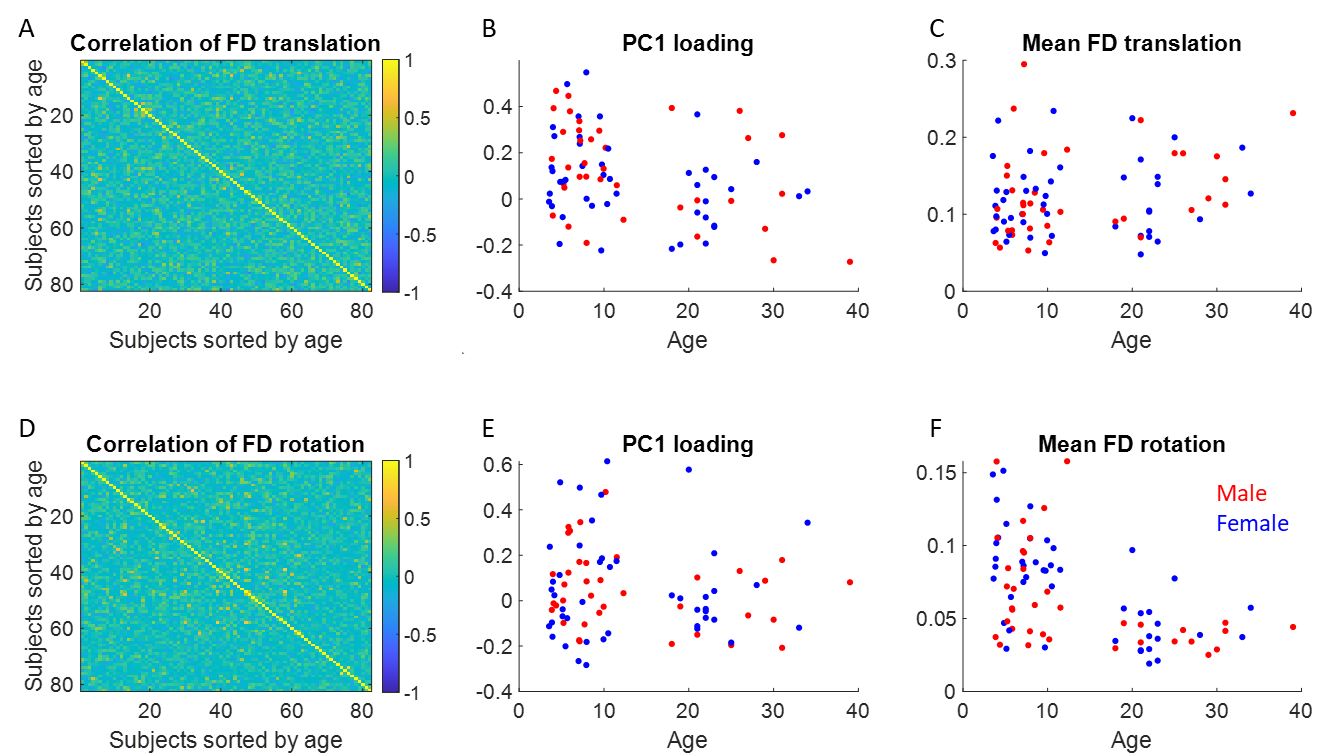

**Figure S1**

Figure S1C and S1F show the mean FD in the two directions, where the mean FD in rotation seemed to have an age effect. We performed 2-sample t-test between the children and adult groups, and confirmed that there was no significant group differences in translation (*t = 1.04, p = 0.30*), but a significant difference in rotation (*t = 6.04, p < 0.001*). Because of the group differences in mean FD, for the analysis of age effects or behavioral correlations, we regressed out the mean FD from PC loadings, can compared the effects before and after regression.

For the supramarginal network (IC17) from the ICA-based analysis, we calculated the correlations between the first two PC loadings and the individual’s head movements. As shown in Table S1, mean FD in rotation showed small but statistically significant correlations with both PC1 loading (*r = -0.30, p = 0.007*) and PC2 loading (*r = 0.29, p = 0.009*). We therefore regressed out the mean FD in the two directions from the two PC loadings. The age effects remain very similar to what before head movement regression (comparing Figure S2 to Figure 3 in the main text).

| r(p) | PC1 loading | PC2 loading | Theory of mind performance |
| --- | --- | --- | --- |
| Mean FD (translation) | 0.06 (0.579) | -0.16 (0.161) | 0.16 (0.250) |
| Mean FD (rotation) | **-0.30 (0.007)** | **0.29 (0.009)** | -0.10 (0.467) |

**Table S1**

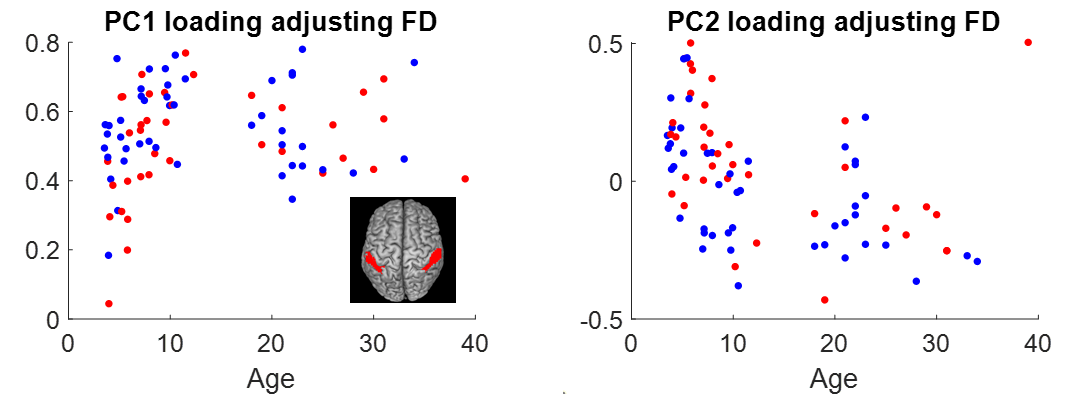

**Figure S2**

The theory of mind performance was not correlated with the mean FD in either directions (Table S1). We regressed out the mean FD in both translation and rotation from the PC loadings. The correlations between adjusted PC loadings and the theory of mind performances remained significant (PC1: *r = 0.428, p = 0.001*; PC2: *r = -0.278, p = 0.044*).

For the first two PC loadings from the precuneus and left sensorimotor region, we also regressed out the mean FD from the two directions. The age effects of the PC loadings were also very similar to what before head motion regression (comparing Figure S3 to Figure 7 and 8 in the main text).

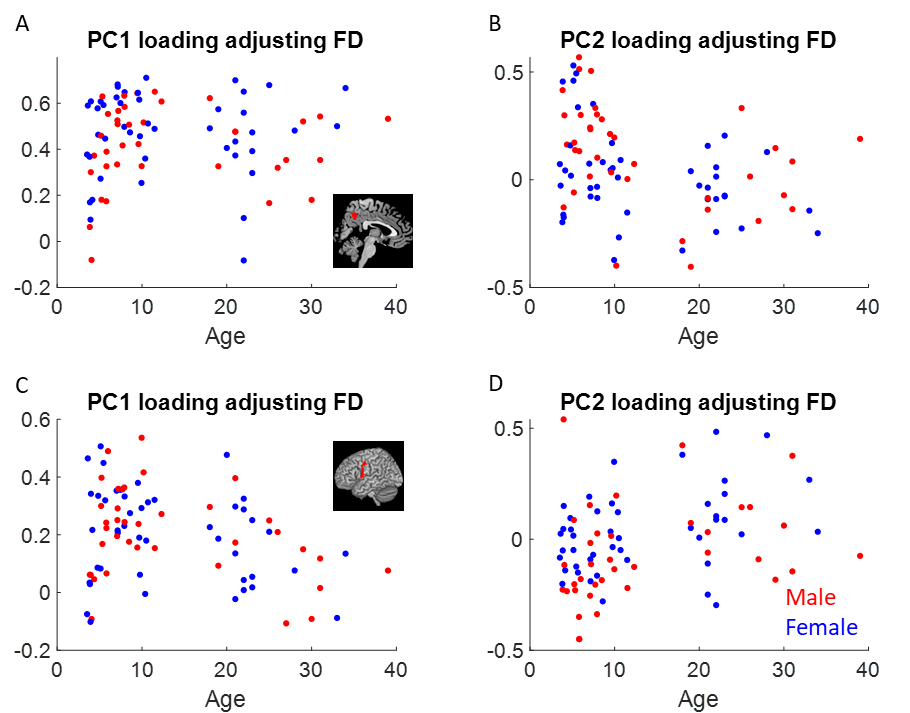

**Figure S3**

**S2. Miscellaneous Analysis on the second PC of the supramarginal network**

Figure S4 shows the percentage variance explained by all the principal components in the supramarginal network (independent component, IC 17). The scree plot supports the inclusion of the second PC in the analysis.

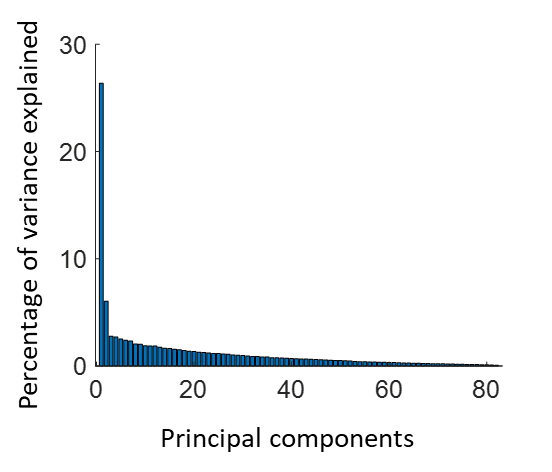

**Figure S4**

To evaluate the stability of the PC2, we performed a bootstrapping analysis on the principal component analysis (PCA) results. Bootstrapping was performed on the subject dimension for 1,000 times. The top row in Figure S5 shows the results for the supramarginal network (independent component, IC 17), where the second PC explained above chance-level variances. Panel A shows the absolute correlation matrix among the first four PCs (PC scores) from the 1,000 bootstrapping samples. The first two PCs were highly correlated among the bootstrapping samples, respectively, indicating that they can be reliably detected. This is in contrast with PC3 and PC4. The bottom row shows the results from the posterior visual network (IC 20), where the second PC was not significant. Panel D shows that the PC2 among the bootstrapping samples did not show reliable correlations among the samples. Panels B and E show the distributions of absolute correlations between PC2 from the original sample and those in the bootstrapping samples, which confirm that the PC2 of supramarginal network was reliable to subject resampling, but that of posterior visual network was not. Panels C and F show the distributions of the percentage variance explained by the second PCs in the bootstrapping samples.

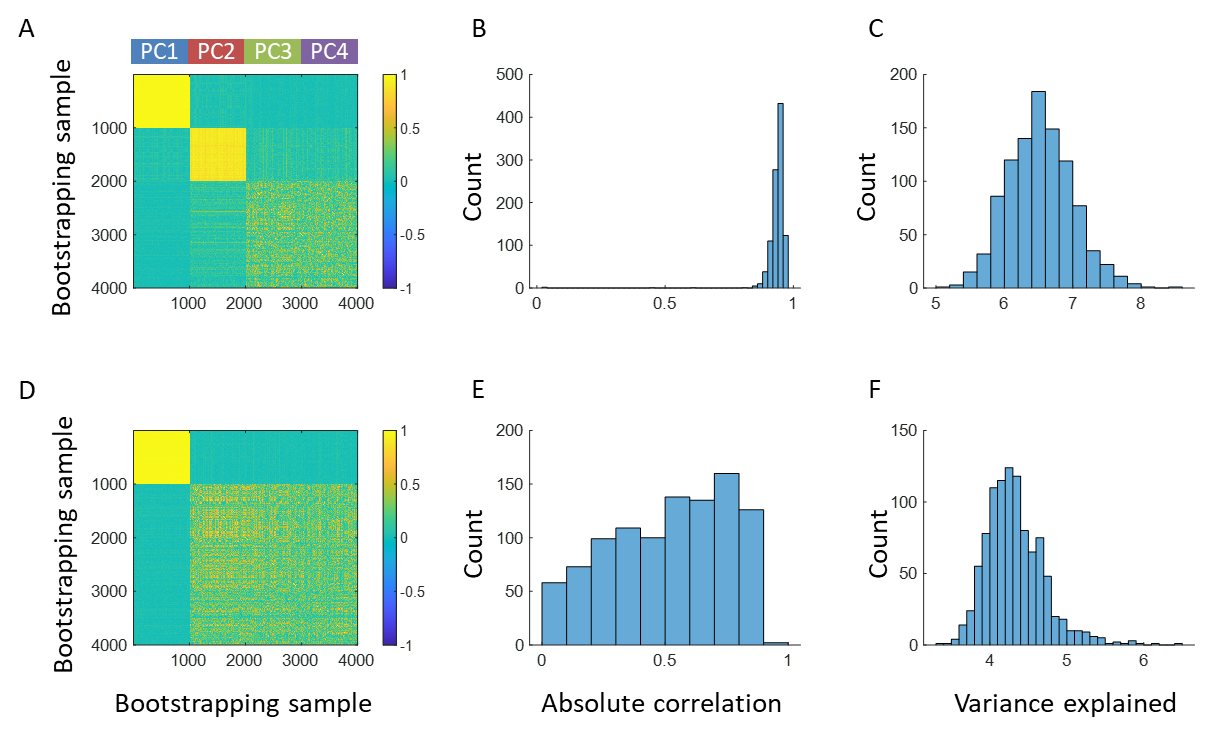

**Figure S5**

**S3. Correlations between PC1 loadings and LOO correlations in the supramarginal network**

Scatter plot between the PC1 loadings and individual leave-one-out (LOO) correlations in the supramarginal network (independent component 17). The red line represents *y = x*.

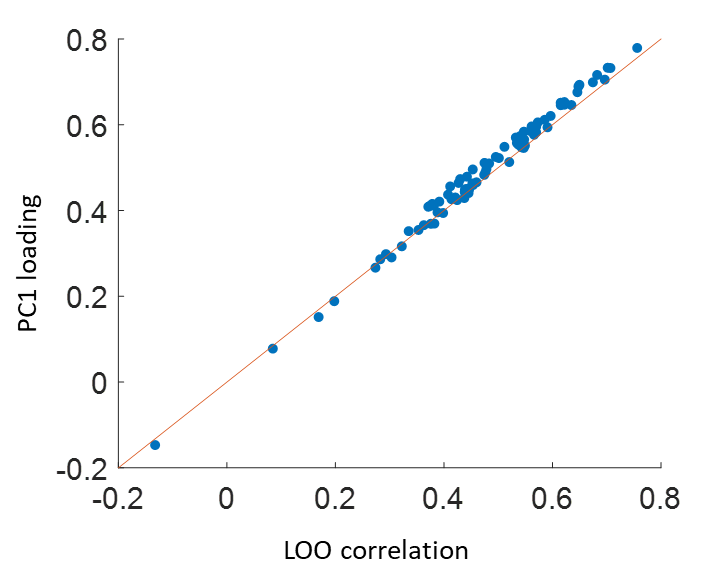

**Figure S6**

**S4. Miscellaneous Analyses on the delay effects in the supramarginal network**

Figure S7 shows the first two PCs in the supramarginal gyrus network (independent component 17), where the two PCs were re-aligned to remove the estimated 2-TR delays.

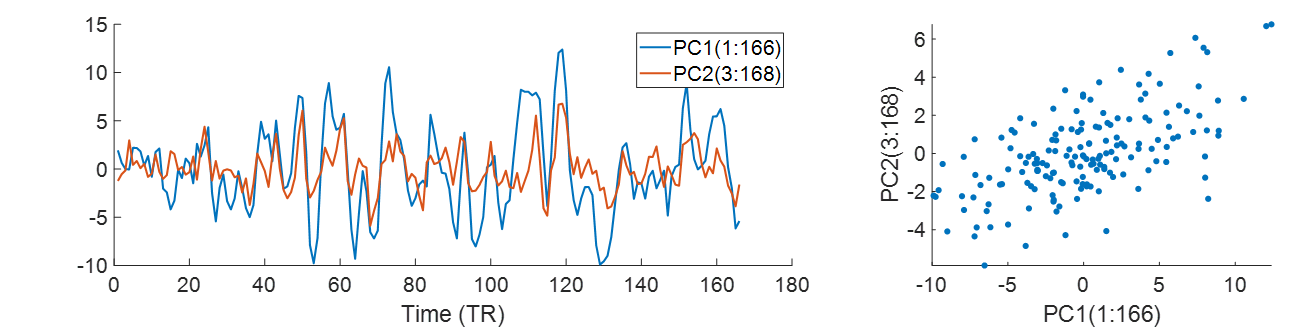

**Figure S7**

Figure S8 shows the fMRI time series in the supramarginal gyrus network (independent component 17) from the 82 subjects sorted according to their age in an ascending order (top) and to the PC2 loadings in a descending order (bottom).

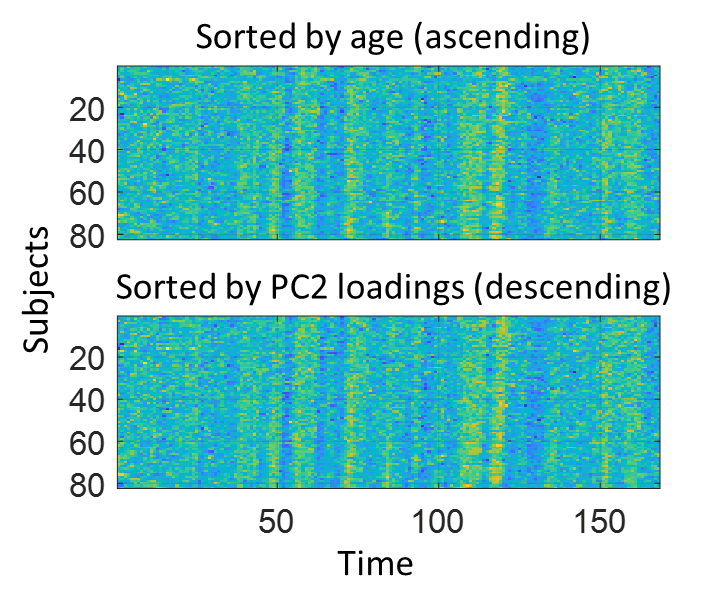

**Figure S8**

**S5. Relations between principal component analysis and LOO correlations**

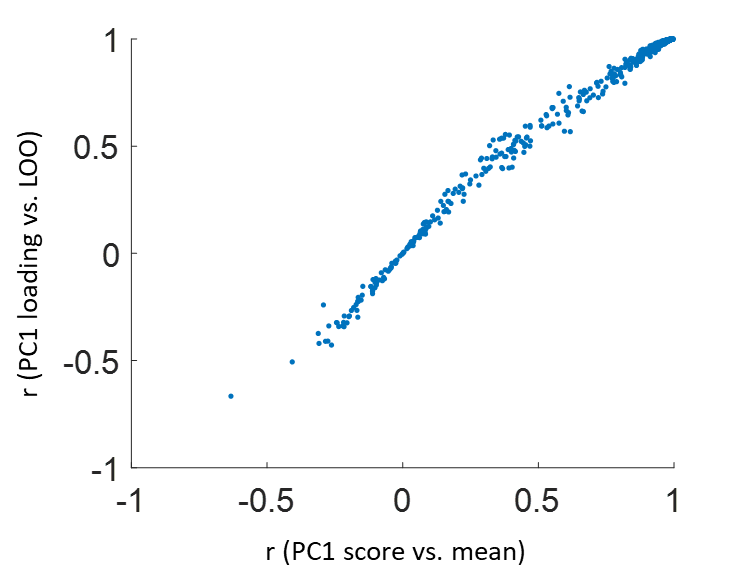

**Figure S9**
